# A Biotin Targeting Chimera (BioTAC) System to Map Small Molecule Interactomes *in situ*

**DOI:** 10.1101/2023.08.21.554211

**Authors:** Andrew J. Tao, Jiewei Jiang, Gillian E. Gadbois, Pavitra Goyal, Bridget T. Boyle, Elizabeth J. Mumby, Samuel A Myers, Justin G. English, Fleur M. Ferguson

## Abstract

Unbiased chemical biology strategies for direct readout of protein interactome remodelling by small molecules provide advantages over target-focused approaches, including the ability to detect previously unknown targets, and the inclusion of chemical off-compete controls leading to high-confidence identifications. We describe the BioTAC system, a small-molecule guided proximity labelling platform, to rapidly identify both direct and complexed small molecule binding proteins. The BioTAC system overcomes a limitation of current approaches, and supports identification of both inhibitor bound and molecular glue bound complexes.

## INTRODUCTION

Small molecule targeted therapies are a cornerstone of modern medicine, providing effective treatments for a wide range of diseases. However, the clinical effects of candidate therapies in patients can be highly variable, even when they bind the same therapeutic target with equal affinity.^1^ This divergent pharmacology may be due to uncharacterized off-target effects, combinatorial polypharmacology, or from small molecules inducing changes in the interactome of their protein targets through allosteric effects or ‘molecular glue’ effects.^2^ Despite the critical role of small molecules in drug discovery and development, there is a lack of comprehensive, network-scale profiling methods that inform on the cellular interactomes of small molecules.

Examples of small-molecule mediated interactome rewiring are well studied in cancer, where targeted therapies frequently induce functional changes in the complexation of their protein targets. For example, blockbuster cancer drugs, such as trametinib^1^ and lenalidomide^3-5^, exert efficacy through the promotion of novel protein complexes, now known as ‘molecular glue’ pharmacology. Unanticipated protein complex rewiring is also a major cause of drug candidate failures, for example underpinning adverse effects of 1^st^ generation RAF inhibitors in RAS driven tumors^6^, and resistance to BET bromodomain inhibitors in triple-negative breast cancer and neuroblastoma^7,8^. However, the discovery of interactome changes that mediate both drug efficacy and drug resistance has so far been serendipitous, as researchers investigate why certain drugs display unexpected pharmacology in the clinic. Despite their central importance, effects on target complexation remain uncharacterized for most protein ligands, representing a ‘blind spot’ in compound characterization workflows.

Current gold-standard technologies for unbiased target-ID are the chemical proteomic techniques photoaffinity labelling^9^ and microenvironment mapping (μMap)^10^ that use UV-light initiated diazirine photochemistry to label liganded proteins with affinity handles. Here, off-compete experiments with free drug, or comparison to a chemically matched negative control molecule facilitate data interpretation. However, in the intracellular context these techniques are directed towards detection of the primary target(s) only, due to the short half-life of the generated reactive carbene species which corresponds to a labelling radius of ∼ 6 nm. Here, the linker length between the diazirine or iridium photocatalyst and the drug dictates the labelling radius, and is therefore limited by cell permeability of the conjugate.^10^ As such, they have yet to be applied to map drug-bound complexes. Diazirine photochemistry approaches are also currently incompatible with *in vivo* applications.

To understand drug-induced interactome changes, affinity purification coupled to mass spectrometry (AP/MS) and proximity labeling coupled to mass spectrometry are commonly employed.^11^ Proximity labeling techniques such as BioID and TurboID are particularly advantageous in mapping interactomes. Proximity labeling methods have a labeling radius of up to 35 nm and can be used in live cells and animals.^12^ By fusing a target protein or localization tag with a proximity labeling enzyme proximity labeling can reliably detect transient, moderate affinity protein-complex interaction *in situ* due to the ability of biotin ligase to accumulate affinity tags on these protein partners over time.^13^ However, both techniques rely on *a priori* target knowledge, which is often incompletely understood for drug candidates, and the fusion of the target protein to either an affinity-tag or a proximity labeling enzyme, which can significantly impact interacting proteins.^13^ Therefore, they cannot be used for unbiased interactome-ID of small molecules. The outputs of these techniques are also typically large numbers of enriched proteins, due to their low stringency, making data interpretation and validation challenging.

To facilitate routine evaluation of ligand-target interactome changes induced by either inhibitors or molecular glues in a single experiment, we envisioned a method that combines the precision and unbiased nature of chemical proteomics with the sensitivity, whole-organism compatibility, and extended detection radius of proximity-labelling coupled to mass spectrometry. Here, we report the development of a ligand-guided miniTurboID method to accomplish these goals, and benchmark it against gold-standard unbiased technologies for both target-ID and interactome-ID.

## RESULTS

Our method, named the biotin targeting chimera (BioTAC) system, uses bifunctional molecules composed of a compound-of-interest linked to selective FKBP12^F36V^ recruiter orthoAP1867, to recruit a ligandable proximity labeling enzyme (mTurbo-FKBP12^F36V^) to compound-bound complexes enabling their biotinylation and subsequent affinity purification (Figure 1a-b). To benchmark the BioTAC system for accurate detection of the direct targets of ligands we selected the well-characterized BET protein inhibitor (+)-JQ1, which potently binds BRD2, BRD3 and BRD4 (as well as the testes-specific BET protein BRDT), as a test case.^14^ We synthesized a series of bifunctional molecules consisting of (+)-JQ1 conjugated to orthoAP1687 via a variable linker (Figure 1b, Figure S1a-c). We measured (+)-JQ1-bifunctional molecule cell permeability using an FKBP12^F36V^ cellular target engagement assay adapted from Nabet *et. al*. Briefly, FKBP12^F36V^ binding molecules compete with an FKBP12^F36V^ targeting PROTAC (dTAG-13), for NLuc-FKBP12^F36V^ occupancy, rescuing degradation and resulting in an increase in NLuc/FLuc (control) signal relative to DMSO-treated cells.^15^ All synthesized compounds were comparably cell permeable (Figure S1d-e)^15^.

**Figure 1.**
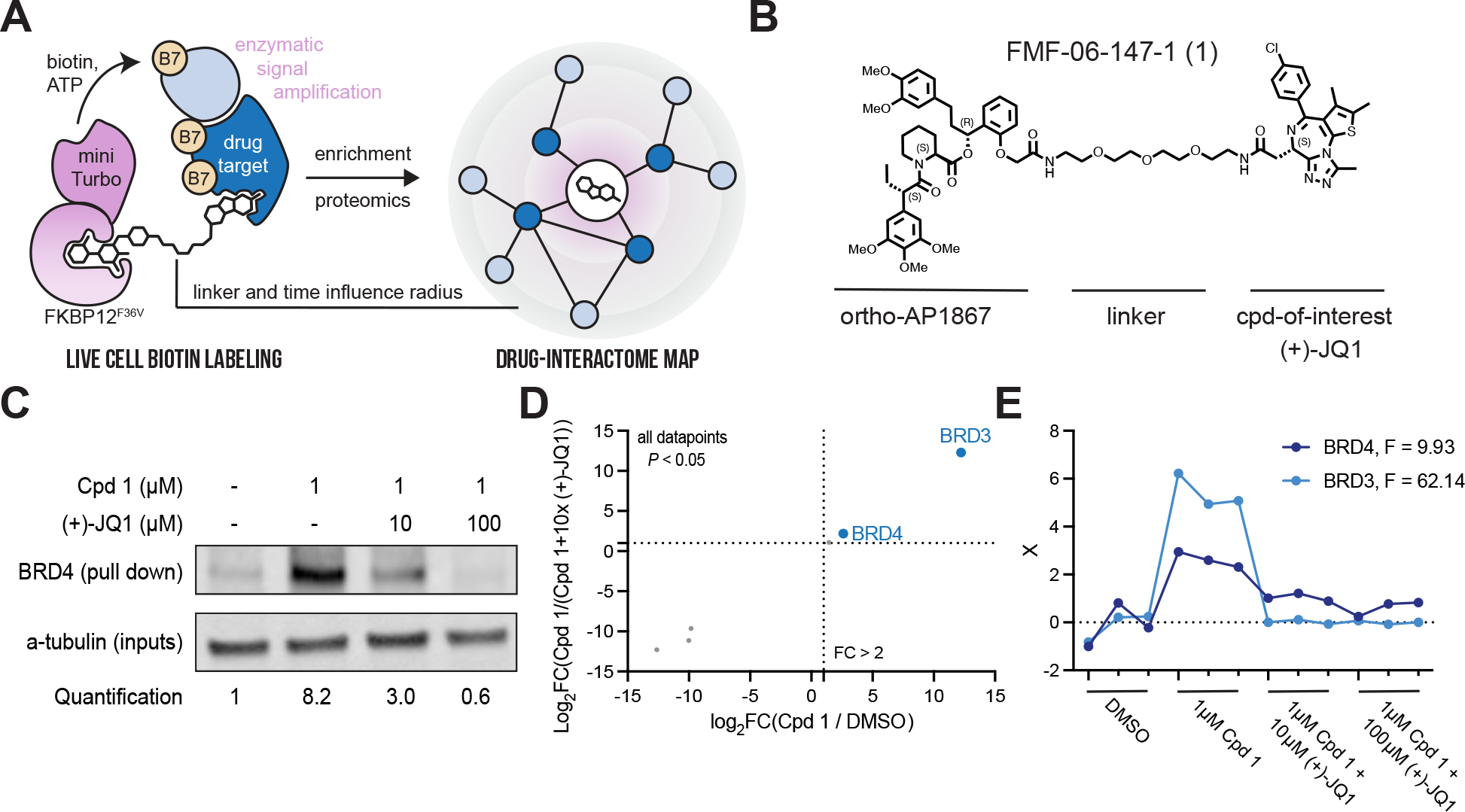
The BioTAC system enables rapid, accurate small molecule target-ID. A. Schematic depicting the components of the BioTAC system. B. Example molecule, Cpd 1, annotated with key functional groups. C. Immunoblot analysis of BRD4 enrichment following treatment of HEK293 cells transiently transfected with miniTurboFKBP12^F36V^ with the indicated compounds and 100 μM biotin at the 30 min timepoint. Data representative of *n* = 2 biologically independent experiments (SI Figure 1J). D. Scatterplot displaying relative FC of streptavidin-enriched protein abundance following treatment of HEK293 cells transiently transfected with miniTurboFKBP12^F36V^ with the indicated compounds and 100 μM biotin at the 30 min timepoint. Only proteins with *P*-value < 0.05 in both conditions depicted. Complete datasets in Table S1, plotted individually Figure S3. E. F-test analysis of proteomic data depicted in D., showing significant enrichment of BRD3 and BRD4 in the presence of 1 μM Cpd 1, relative to DMSO and (+)-JQ1 off-compete experiments. F-statistic listed top right, X = scaled abundance ratio.

We next performed bifunctional molecule guided-proximity labeling experiments using our validated reagents. HEK293 cells transiently transfected with mTurbo-FKBP12^F36V^ were treated with 100 μM biotin, 1 μM bifunctional (+)-JQ1 recruiters, and variable concentrations of free (+)-JQ1 to off-compete the bifunctional molecule and rescue biotinylation. Biotinylated proteins were isolated from cell lysates via streptavidin bead pulldown and analyzed by western blot for BRD4, a primary target of (+)-JQ1. We observed significant enrichment of BRD4 and a dose dependent decrease in BRD4 pulldown in the presence of unconjugated (+)-JQ1 at all timepoints evaluated (Figure 1c, Figure S1g-j). Comparable activity was observed for all bifunctional analogues, indicating low sensitivity to linker length (Figure S1g-j), FMF-01-147-1 (Cpd 1) was selected for further characterization based on high BRD4-labeling and rescue at the 30 min time point (Fig. 1c).

Having determined that the BioTAC system could identify BRD4 as a (+)-JQ1 target in focused screens, we sought to evaluate it as an unbiased target-ID method. To identify direct binders of (+)-JQ1 proteome-wide, we performed BioTAC proximity labelling experiments as described above, followed by label-free mass spectrometry-based proteomic analysis of biotinylated proteins enriched following a 30 minute treatment with 100 μM biotin plus DMSO or 1 μM Cpd 1, and then 1 μM Cpd 1 off-competed by pre-treatment with 10 μM free (+)-JQ1. We observed highly selective enrichment of known (+)-JQ1 targets BRD3 and BRD4, comparable to published (+)-JQ1 μMapping and superior to published (+)-JQ1 photoaffinity labelling (Figure 1d-e, Figure S2a-c, Figure S3a-c, Table S1).

Having established conditions for determining the primary targets of small molecules using the BioTAC system, we next investigated its utility in reading out the interactome sphere of (+)-JQ1 bound BET-proteins. An advantage of the BioTAC system is the relatively long half-life of the activated biotin-AMP intermediate generated by TurboID, allowing the labelling radius to be increased up to 35 nm by extending the labelling time.^12^ To quantitively evaluate the ability of the BioTAC system to successfully enrich the known interactome of (+)-JQ1 bound BET proteins, we performed time course BioTAC proximity labelling experiments at the 1 h and 4 h time points, and evaluated streptavidin-enriched biotinylated proteins using mass spectrometry-based proteomic analysis (Figure 2a-c, Figure S3, Table S1). As expected, longer time points correlated with a greater number of proteins meeting our significance cut-offs (defined as FC > 2, *P* < 0.05) for both enrichment in Cpd 1 vs. DMSO, and competition with free (+)-JQ1 in Cpd 1 vs. Cpd 1 + 10 μM (+)-JQ1, (Figure 2a, Figure S3). Known (+)-JQ1 direct targets BRD3 and BRD4 were significantly enriched and off-competed at all time points and BRD2 was also detected at the 4 hr timepoint. Encouragingly, identified hit proteins at 1-4 hrs also included many known BET interactors, such as proteins found in components of the Pol II productive elongation complex subunit P-TEFb, the TFIID complex, and the Nucleosome Remodeling Deacetylase (NuRD) complex, as well as known BRD4 direct-binder Histone H4 (Table S1). These observations suggested that we may be labelling direct interactors of (+)-JQ1 bound BRD2, 3, and 4. To test this hypothesis we compared our hits to an extensive BET protein interactome reference dataset identified using AP/MS and proximity labelling coupled to mass spectrometry with and without (+)-JQ1, reported by Lambert *et al*.^16^ The BioTAC system afforded significant enrichment and off-competition (*P* < 0.0001, one-way ANOVA) of identified reference interactors at the 4 h time point (Figure 2b). We next asked whether we could perform the inverse, enriching complex members identified by Lambert *et al*. directly from our data without any *a priori* knowledge of known interactors. We were pleased to discover that 60% of our statistically significant enriched and (+)-JQ1 rescued hits were also present in the Lambert dataset. This observation falls greater than 7 standard deviations above a random bootstrap analysis of our data without filtering for significant enrichment and (+)-JQ1 rescue (Figure 2c). To account for the limitations of using the findings of only one study as a benchmark, we performed Gene Ontology (GO) Biological Process analysis of the hits from the 4hr timepoint^17,18^. Here, transcriptional elongation by Pol II was significantly enriched (P < 0.05), consistent with the known functions of BET protein containing complexes targeted by bromodomain inhibitors (Figure 2d).^19^

**Figure 2.**
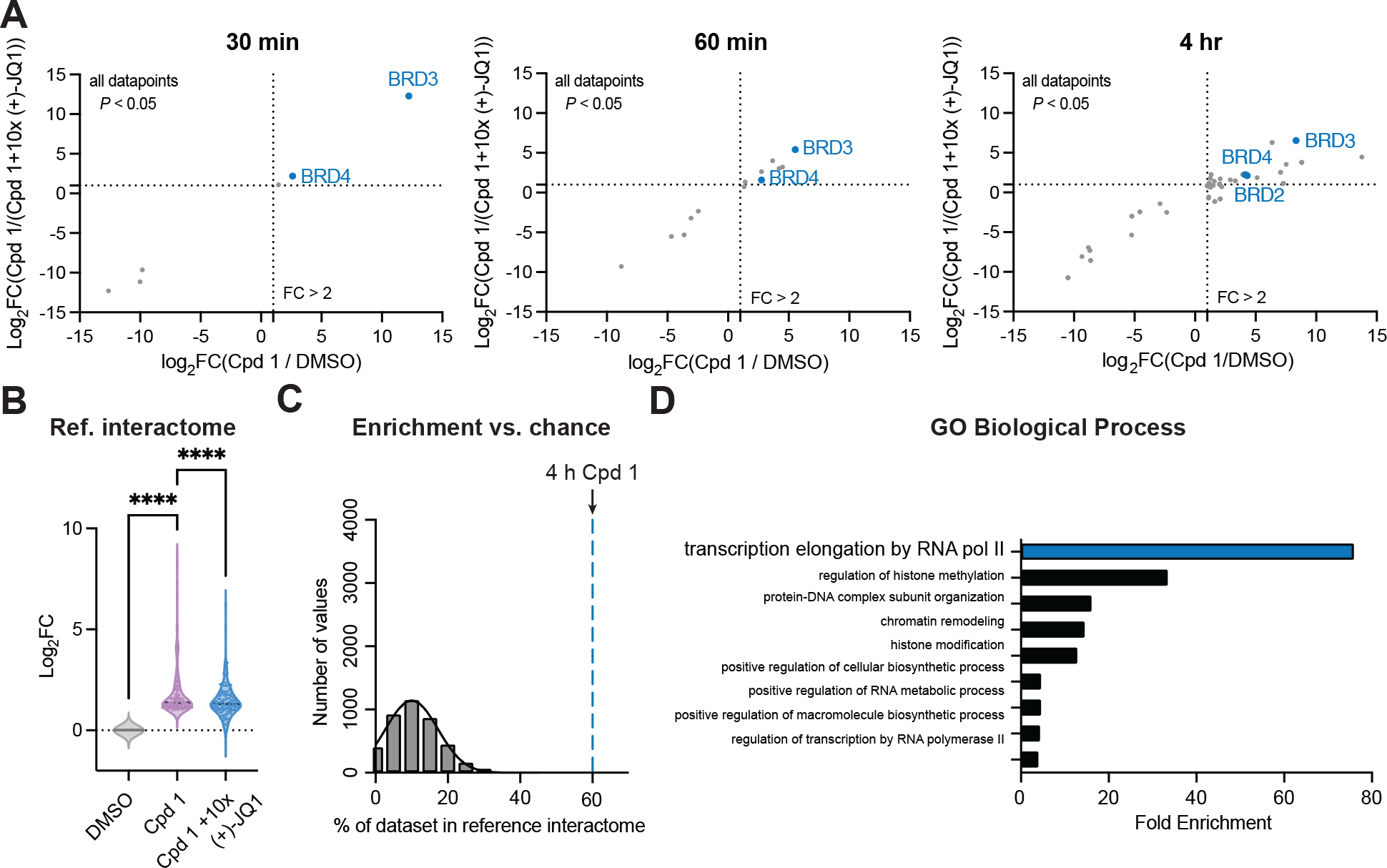
The BioTAC system enables rapid, accurate small molecule interactome-ID. A. Time-course proteomics of streptavidin-enriched biotinylated proteins isolated from HEK293 cells transiently transfected with miniTurboFKBP12^F36V^ and treated with the indicated compounds and 100 μM biotin, demonstrating enrichment and competition of known direct targets (30 min) and complexed proteins (60 min, 4 hrs). Only proteins with *P*-value < 0.05 in both conditions depicted. Complete datasets in Table S1, plotted individually Figure S3. High-confidence hits are defined as those that are enriched > 2-fold in both Cpd 1/ DMSO and Cpd 1 / Cpd 1 + 10x (+)-JQ1, where *P* < 0.05, plotted upper-right quadrant. B. Fold-change of all reference BRD2, BRD3, and BRD4 interactors (Lambert *et. al*.) enriched in the Cpd 1 vs. DMSO dataset at the 4 hr time point, showing statistically significant rescue (*P* < 0.0001, 1-way ANOVA). C. Percent enrichment of reference interactors (Lambert *et. al*.) in the hits identified at the 4 hr timepoint (dotted line), vs. the percent enrichment of known interactors in 4,000 random protein sets of equivalent size from the DMSO control (gray bars), showing significant enrichment by Cpd 1 BioTAC (> 7 standard deviations away from mean of random chance). D. Gene Ontology analysis, showing significant enrichment of biological processes associated with known BET-protein function in the 4 hr Cpd 1 BioTAC experiment hits.

The paucity of practical methods for rapid, unbiased readout of small-molecule induced interactome changes has hindered the rational discovery and development of molecular glues. Molecular glues are small molecules which exert their function by to binding a primary target, and promoting or strengthening its interaction with a second protein through interactions at the protein-protein interface.^20^ Molecular glue discovery is an area of high biomedical interest due to the ability of molecular glues to target undruggable oncoproteins such as transcription factors, which are recalcitrant to traditional inhibitor discovery but can be neutralized by induced complexation and targeted degradation.^20^ However, as the binary affinity between a molecular glue and the second recruited target is low or non-existent in the absence of the primary target, molecular glue interactions are challenging to detect and screen for. Having rigorously benchmarked the performance of the BioTAC system using well-characterized (+)-JQ1, we sought to use the BioTAC system to inform on complexes assembled by molecular glues (Figure 3a).^20^ We selected Trametinib as a non-dergrader glue with which to benchmark the platform. Trametinib derives its clinical anti-cancer efficacy from promoting the interaction of its primary target MEK1/2 with KSR1, but has low affinity for KSR1 alone meaning this interaction was missed during clinical developments.^1^

**Figure 3.**
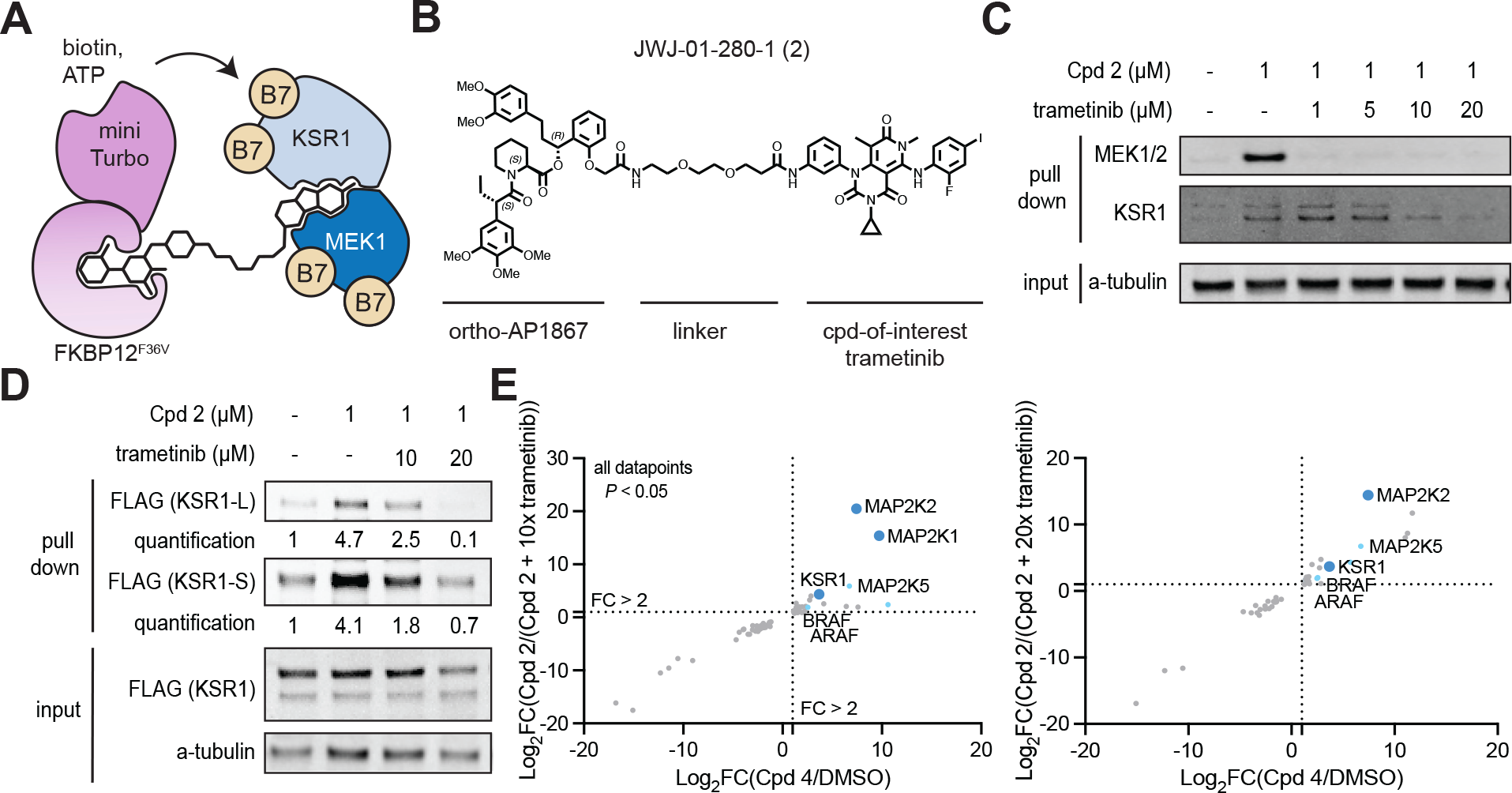
The BioTAC system enables detection of non-degrader molecular glue interactions. A. Schematic depicting how the BioTAC system can detect molecular glue interactions. B. Chemical structure of Trametinib recruiting bifunctional molecule JWJ-01-280-1/Cpd 2. C. Immunoblot analysis of MEK1 and KSR1 following treatment of HEK293 cells transiently transfected with miniTurboFKBP12^F36V^ with the indicated compounds and 100 μM biotin at the 4 h timepoint, and streptavidin-based enrichment, showing successful enrichment and competition with trametinib. D. Immunoblot analysis of mKSR1 following treatment of HEK293 cells transiently transfected with miniTurbo-FKBP12^F36V^ and mKSR1, with the indicated compounds and 100 μM biotin at the 4 h timepoint, and streptavidin-based enrichment, showing successful enrichment and competition with trametinib. In C, D two KSR1 isoforms are observed, produced by alternative splicing.^26^ KSR1-L (102 KDa) corresponds to the expected MW of Uniprot Q8IVT5-1 (canonical sequence), KSR1-S (87 KDa) corresponds to the expected MW of variant with residues 1-137 missing, Uniprot Q8IVT5-3, and -4. Our data indicate both isoforms can complex with trametinib-bound MEK1.

We synthesized bifunctional trametinib analogue JWJ-01-280/Cpd 2, with a linker attachment informed by a reported trametinib-derived BODIPY-linked BRET probe named Tram-bo (Figure 3b).^1^ We evaluated the cell permeability of Cpd 2 as described above, which was less cell permeable than the (+)-JQ1 derivatives (Figure S4a). Nevertheless, Cpd 2 supported efficient MEK1 labelling, following dose and time point optimization (Figure S4b-c). We evaluated the ability of the BioTAC system to detect both known interactors of trametinib by Western blot, as described above. We successfully detected the MEK1:trametinib:KSR1 complex using the BioTAC system following a 4 hr treatment with 1 μM Cpd 2, and dose-dependent competition in the presence of trametinib of both KSR1 and MEK1 by immunoblot (Figure 3c, Figure S6). Low expression of KSR1 in HEK293 cells resulted in lower KSR1 signal relative to MEK1 in both input and enriched samples, indicating that a small proportion of MEK1 in HEK293s is bound by KSR1 in the presence of trametinib. In line with the reported finding that trametinib binds more tightly, and with a slower off-rate (K_dis_) to MEK1-KSR1 than to MEK1 alone, higher concentrations of trametinib were required to off-compete labelling of KSR1 than MEK1.^1^ To strengthen our confidence that the MEK1: trametinib: KSR1 complex can be reliably detected by BioTAC, and to allow quantification of enrichment and competition, we turned to published conditions from Khan *et al*.^1^ We repeated the BioTAC experiment, this time with mKSR1 overexpressed concomitantly with mTurbo-FKBP12^F36V^, and observed significant enrichment of KSR1 in the presence of Cpd 2 and convincing off-competition by trametinib (Figure 3d, Figure S6). To examine the global interactome of Trametinib in cells, we performed BioTAC experiments with Cpd 2, coupled to mass spectrometry. Here, we successfully enriched KSR1, alongside other known MEK1/2 interactors AFAR and BRAF (Figure 3e). Together, these data demonstrate the ability of the BioTAC system to detect the complexes assembled by non-degrader molecular glues.

## DISCUSSION

Methods for the routine measurement of drug-target interactomes are lacking, hiding the mechanism of action of numerous small molecules from view. Even though interactome remodelling in response to small molecule drugs is a common phenomenon that mediates drug efficacy and resistance, little progress has been made in identifying and characterizing such events. Here we report the BioTAC system, which can identify the direct target of a small molecule, as well as its complexed proteins with high confidence. We demonstrate successful enrichment of the reported interactomes of the epigenetic inhibitor (+)-JQ1 and the molecular glue trametinib, but this approach is theoretically applicable to any small molecule of interest that can be functionalized with a linker.

Next, we show the BioTAC system can identify molecular glue pairs that previously evaded detection, in a single experiment using trametinib, MEK1/2, KSR1, as a model system. The discovery and detection of molecular glue interactions is notoriously challenging to evaluate, in particular when evaluating non-degrader molecular glues. Recently, elegant workflows consisting of mechanism-based screening in wild-type and hypo-NEDDylated cells, followed by multi-omic target deconvolution have been described for the discovery of cullin-ring ligase (CRL)-recruiting molecular glue degraders.^21,22^ To identify non-degrader molecular glues, size exclusion chromatography coupled to mass spectrometry has been used in combination with activity-based protein profiling to screen electrophilic compound libraries, yielding covalent stabilizers and disruptors of protein-protein interactions.^23^ However, these approaches require resource intensive multi-omic workflows, and are limited to covalent glues, or CRL-mediated degradation mechanisms. In future applications, we envision the BioTAC system may be used in a screening mode, for unbiased profiling of putative glue libraries.

The BioTAC system uses a universal recruitable biotin ligase chimera, miniTurbo, facilitating rapid application. Bifunctional molecules for investigating any drug-of-interest are synthesised in one step from a common ortho-AP1867 precursor using robust coupling chemistries. Finally, the enrichment, mass spectrometry and data analysis methods are adapted from standard protocols in proximity labelling already performed by most proteomics core facilities.^24^ These features make BioTAC system readily accessible for broad application. During the preparation of this manuscript, a preprint describing a related approach for identifying targets of small molecules via SNAP- and Halo-tagging of TurboID was also disclosed.^25^ Whilst these systems differ from the BioTAC system in their reported specificity, and have not yet been benchmarked for interactome-detection, their successful implementation across a range of ligands highlights the robustness of using proximity labelling to interrogate small molecule targets.^25^

In the long term, building community-wide knowledge around how small molecule drugs alter their target proteins complexation will lay the foundation for the rational design of drug target interactome profiles, to combat drug resistance, and enable wider targeting of the undruggable proteome.

## AUTHOR CONTRIBUTIONS

F.M.F. and J.G.E. conceived and led the study. J.G.E. designed and cloned the DNA constructs. J.W.J. and F.M.F. designed and performed molecule synthesis. A.J.T. and G.E.G. performed dual luciferase target engagement assays. A.J.T. and B.T.B. performed immunoblot experiments. A.J.T. prepared samples for proteomic studies. A.J.T., S.A.M. and J.G.E. performed proteomic data analysis. F.M.F. wrote the manuscript with input from all authors.

## ACKNOWLEDGEMENTS

This work was supported by American Cancer Society IRG Grant # IRG-19-230-48-IRG, UC San Diego Moores Cancer Center, Specialized Cancer Center Support Grant NIH/NCI P30CA023100, and NIH grant DP2NS132610 (F.M.F.). A.J.T. was supported by a UCSD Distinguished Graduate Student Award. G.E.G. was supported by the Molecular Biophysics Training Grant T32 32GM139795. This study was supported by NIH grant DP2GM146247 (J.G.E.). The authors would like to thank Dr. Majid Ghassemian, Dr. Lisa Jones, and Dr. Katherine A. Donovan for helpful discussions and feedback and Dr. Sonya Neal for use of a BioRad ChemiDoc Gel Imager.

