## Supplementary material for "A Biotin Targeting Chimera (BioTAC) System to Map Small Molecule Interactomes *in situ*": SI Figures 1-4

### Supporting Information

### Contents

Supporting Figure 1 | Optimization of (+)-JQ1 orthoAP1867 bifunctional molecules……………..2

Supporting Figure 2 | Comparison of the BioTAC system to photoaffinity labeling and µMap…..3

Supporting Figure 3 | Volcano plots for all proteomics experiments………………………………..4

Supporting Figure 4 | Optimization of MEK1 labeling by trametinib bifunctional molecules……..5

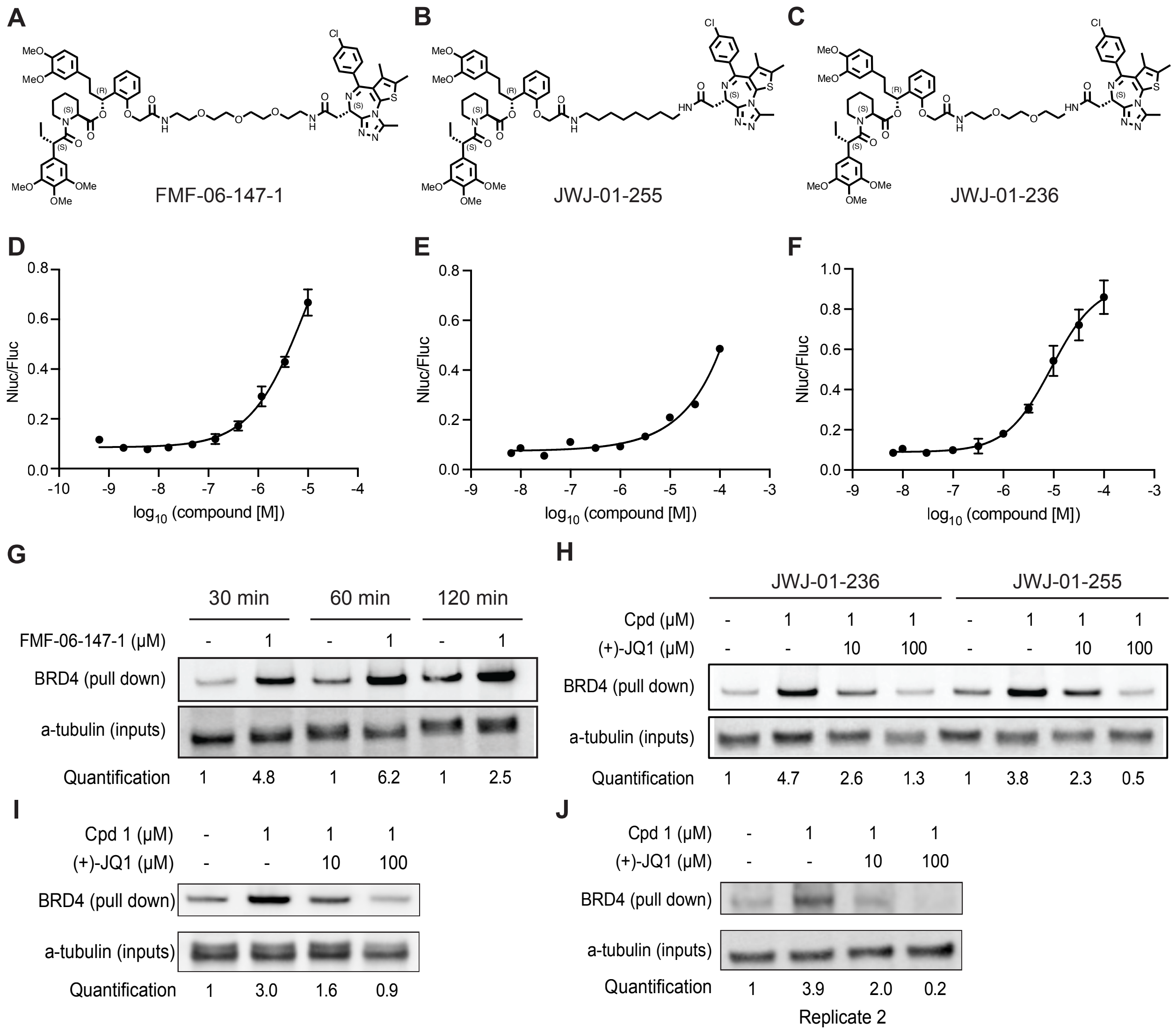


**Supporting Figure 1 | Optimization of (+)-JQ1 orthoAP1867 bifunctional molecules.** A.-C. Chemical structure of all tested (+)-JQ1-linker-ortho-AP1867 analogues. D. FKBP12$^{F36V}$ cellular target engagement assays for compounds shown A.-C., data plotted as mean ± S.D. of n = 3 technical replicates. D. FMF-06-147-1, E. JWJ-01-255, F. JWJ-01-236, demonstrating competitive displacement of dTAG-13. G. Immunoblot analysis of BRD4 enrichment following treatment of HEK293 cells transiently transfected with miniTurboFKBP12$^{F36V}$ and treated with FMF-06-147-1 and 100 μM biotin at the indicated timepoint. H. Immunoblot analysis of BRD4 enrichment following treatment of HEK293 cells transiently transfected with miniTurboFKBP12$^{F36V}$ and treated with the indicated compounds and 100 μM biotin at the 60 min timepoint. I. Immunoblot analysis of BRD4 enrichment following treatment of HEK293 cells transiently transfected with miniTurboFKBP12$^{F36V}$ and treated with FMF-06-147-1 and 100 μM biotin at the 30 min timepoint. J. Replicate 2 of immunoblot analysis of BRD4 enrichment following treatment of HEK293 cells transiently transfected with miniTurboFKBP12$^{F36V}$ and treated with FMF-06-147-1 and 100 μM biotin at the 30 min timepoint. G., I. Data representative of *n* = 3 biologically independent experiments. J. Data representative of *n* = 2 biologically independent experiments. S.D. Standard deviation.

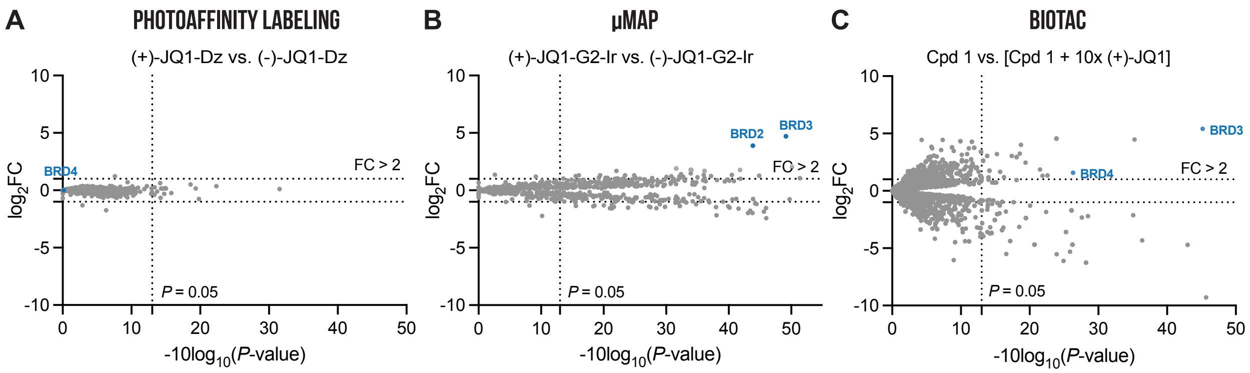


**Supporting Figure 2 | Comparison of the BioTAC system to photoaffinity labeling and μMap.** A.-B. Replotted reference data from Trowbridge *et. al.* showing photoaffinity labeling and μMap results for (+)-JQ1. C. Proteomics analysis of biotinylated proteins enriched by streptavidin pulldown from HEK293 cells transiently transfected with miniTurbo-FKBP12$^{F36V}$ and treated with 100 μM Biotin, 1 μM of Cpd 1 ± 10 μM (+)-JQ1 for 30 mins.

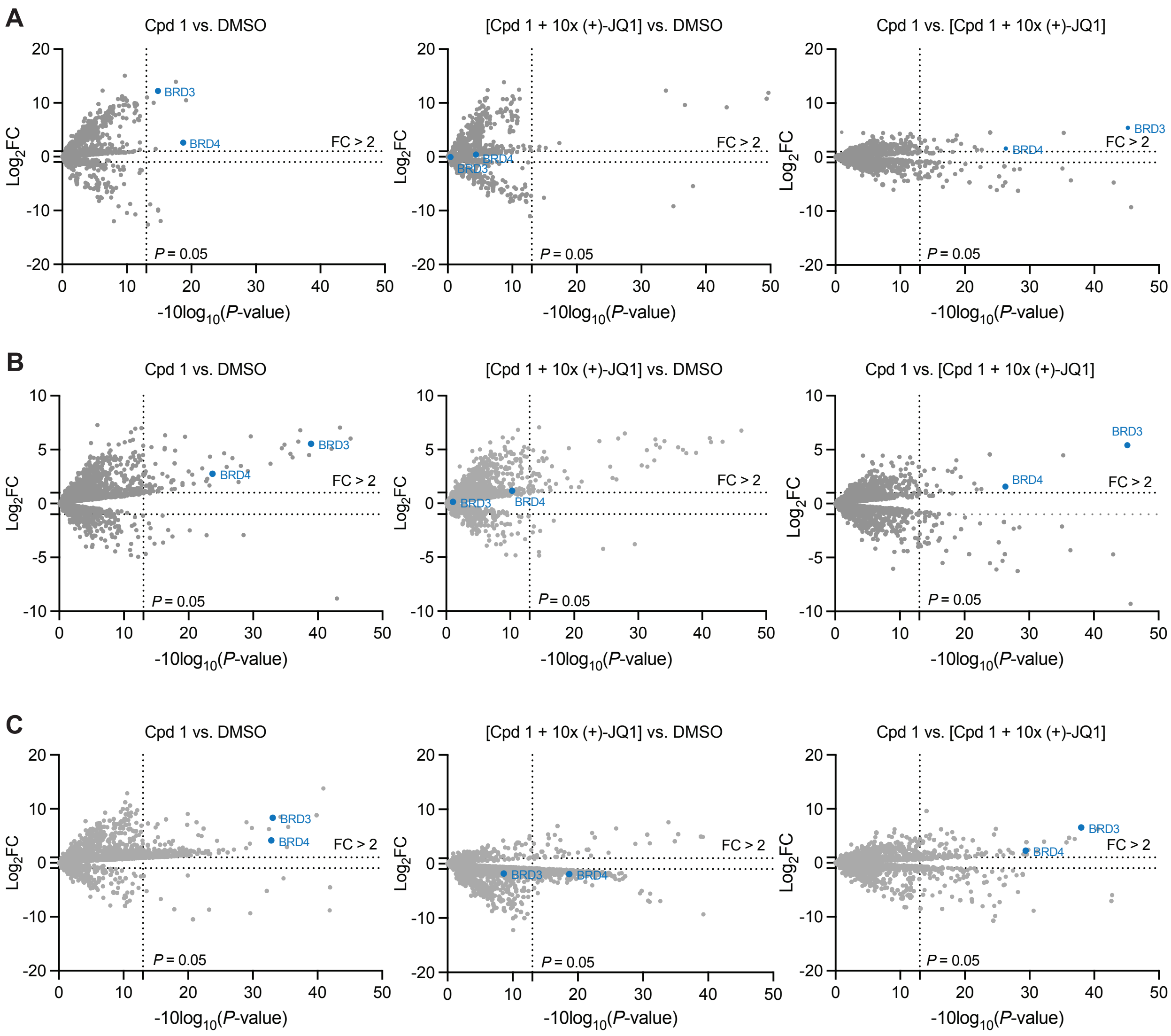


**Supporting Figure 3 | Volcano plots of proteomic experiments.** Volcano plots showing all datapoints from mass-spectrometry based proteomic analysis of proteins enriched by streptavidin pulldown from HEK293 cells transiently transfected with miniTurbo-FKBP12$^{F36V}$ and treated with 100 μM Biotin, DMSO or 1 μM of Cpd 1 ± 10 μM (+)-JQ1 for A. 30 mins. B. 60 mins. C. 4 hrs. Points corresponding to BRD3 and BRD4 are highlighted blue.

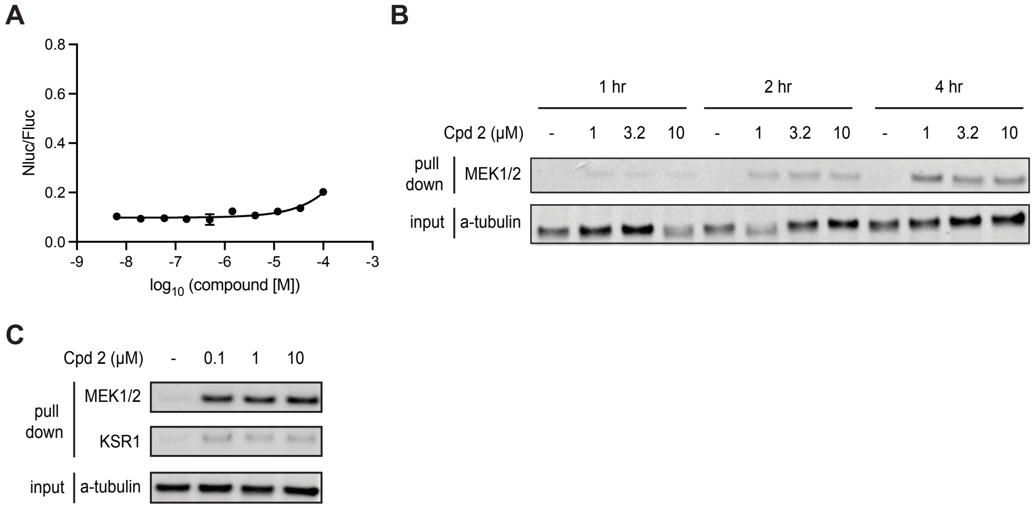


**Supporting Figure 4 | Optimization of MEK1 labeling by trametinib bifunctional molecules.** A. FKBP12$^{F36V}$ cellular target engagement assays for JWJ-01-280, data plotted as mean ± S.D. of n = 3 technical replicates. B. Immunoblot analysis of MEK1/2 enrichment following treatment of HEK293 cells transiently transfected with miniTurboFKBP12$^{F36V}$ with JWJ-01-280-1 and 100 μM biotin at the indicated timepoint. C. Immunoblot analysis of MEK1/2 and KSR1 enrichment following treatment of HEK293 cells transiently transfected with miniTurboFKBP12$^{F36V}$ with the indicated concentration of JWJ-01-280-1 and 100 μM biotin for 4 h. Data representative of *n* = 3 biologically independent experiments. S.D. Standard deviation.
